# SurvivalML: an integrative platform for the discovery and exploration of prognostic models in multi-center cancer cohorts

**DOI:** 10.1101/2022.10.25.513678

**Authors:** Zaoqu Liu, Hui Xu, Long Liu, Siyuan Weng, Zhe Xing, Yuqing Ren, Xiaoyong Ge, Libo Wang, Chunguang Guo, Shuang Chen, Quan Cheng, Peng Luo, Jian Zhang, Xinwei Han

## Abstract

Advances in multi-omics and big-data technologies have led to numerous prognostic signatures aimed at improving current clinicopathological staging systems. Due to the lack of reproducibility and independent confirmation, few signatures have been translated into clinical routine. As high-quality datasets accumulate, identifying robust signatures across multiple independent cohorts becomes possible. Nonetheless, inaccurate data retrieval, different versions of genome annotations, disparate expression distributions, difficult data cleaning, inconsistent clinical information, algorithm selection, and parameter tuning have impeded model development and validation in multi-center datasets. Hence, for the first time, we introduced SurvivalML (https://rookieutopia.com/app_direct/SurvivalML/), a web application for helping develop and validate prognostic models across multi-center datasets. SurvivalML included 37,325 samples (253 eligible datasets) with both transcriptome data and survival information from 21 cancer types, which were renewedly and uniformly re-annotated, normalized, and cleaned. This application provided 10 survival machine-learning algorithms for flexibly training models via tuning essential parameters online and delivered four aspects for model evaluation, including Kaplan-Meier survival analysis, time-dependent ROC, calibration curve, and decision curve analysis. Overall, we believe that SurvivalML can serve as an attractive platform for model discovery from multi-center datasets.

## Introduction

Cancer represents a complex and aggressive disease with high incidence and mortality worldwide^1^. Current management systems of cancer predominantly rely on the anatomical extent of cancer, such as the American Joint Committee on Cancer (AJCC) staging scheme^2^. Nevertheless, clinicopathological staging systems are insufficient to achieve individuation management due to cancer patients usually exhibiting heterogeneous clinical outcomes^3^. Timely intervention for “high-risk” patients identified by the stratification system is essential to improve their prognosis. Hence, the refinement of stratification strategies needs to consider biological traits for optimizing clinical decision-making^4^.

Fueled by multi-omics and big-data technologies, current cancer studies can investigate molecular features in multiple dimensions and the clinical relevance of massive molecular alterations can be efficiently inferred^4–7^. This progress has also spawned a plethora of gene expression signatures that have been applied in various scenarios, such as distinguishing cancer types, identifying cancer grades, assessing drug sensitivity, and predicting prognosis^4,6,8–11^. Relative to individual biomarkers, a multigene panel consisting of significant biomarkers might be a more ideal way to tackle both intra- and inter-tumor heterogeneity in cross-platform cohorts^4,12^. A previous study developed a 70-gene signature for stratifying prognosis in 295 consecutive patients with breast cancer^9^. Castells and colleagues established a scoring system to evaluate the recurrence-free survival of stage II/III colon cancer receiving adjuvant 5-fluorouracil-based chemotherapy^10^. In clinical settings, survival endpoints are crucial outcomes of cancer patients and the primary basis for clinicians to conduct treatment and surveillance^13^. However, although numerous prognostic signatures were developed, few have been introduced into clinical implementation due to a lack of reproducibility and independent validation in multi-center cohorts^4^.

Currently, the continuous accumulation of public datasets creates opportunities to identify robust signatures across multi-center cohorts^7^. Frustratingly, inaccurate data retrieval, different versions of genome annotations, disparate expression distributions, difficult data cleaning, inconsistent clinical information, algorithm selection, and parameter tuning have hindered the development and validation of prognostic models in multiple independent datasets^5,14–16^. In addition, these works are difficult and inconvenient for clinicians and biologists without computational programming skills. Thus, to address these issues, we developed a free online web application termed SurvivalML (Survival Machine-Learning), for helping develop and validate prognostic signatures via various survival machine-learning algorithms across cleaned multi-center datasets. SurvivalML included a total of 37,325 samples (253 datasets) with both transcriptome data and survival information from 21 cancer types, which have been renewedly and uniformly re-annotated, normalized, and cleaned. Ten survival machine-learning algorithms were provided to flexibly develop and validate models via tuning essential parameters online. Four analytical methods including Kaplan-Meier survival analysis, time-dependent ROC, calibration curve, and decision curve analysis were supplied to comprehensively evaluate the model performance. Overall, we believe that SurvivalML might serve as an attractive platform for model discovery from multi-center datasets, which is available at https://rookieutopia.com/app_direct/SurvivalML/.

## Methods and materials

### Data generation

Public databases have stored large-scale data on various cancer types, laying the foundation for multi-center development and validation of survival models. This study mainly generated data from five databases, encompassing The Cancer Genome Atlas Program (TCGA, https://portal.gdc.cancer.gov), Gene Expression Omnibus (GEO, https://www.ncbi.nlm.nih.gov/geo/), International Cancer Genome Consortium (ICGC, https://dcc.icgc.org), Chinese Glioma Genome Atlas (CGGA, http://www.cgga.org.cn/), and ArrayExpress (https://www.ebi.ac.uk/arrayexpress/) (Figure 1A). Cancer datasets were included if the following criteria were met: i) had both transcriptome data and survival information; ii) no less than 20 samples; iii) no less than 2,500 genes after filtering out non-expressed genes across all samples. In total, we collected 261 survival datasets (n =38,934) from 27 cancer types, including adrenocortical carcinoma (ACC), bladder cancer (BLCA), breast cancer (BRCA), cervical squamous cell carcinoma (CESC), cholangiocarcinoma (CHOL), colorectal cancer (CRC), esophageal carcinoma (ESCA), glioblastoma multiforme (GBM), head-neck squamous cell carcinoma (HNSC), kidney renal clear cell carcinoma (KIRC), liver hepatocellular carcinoma (LIHC), lower grade glioma (LGG), lung adenocarcinoma (LUAD), lung squamous cell carcinoma (LUSC), diffuse large B-cell lymphoma (DLBC), mesothelioma (MESO), ovarian cancer (OV), pancreatic adenocarcinoma (PAAD), pheochromocytoma and paraganglioma (PCPG), prostate adenocarcinoma (PRAD), sarcoma (SARC), skin cutaneous melanoma (SKCM), stomach adenocarcinoma (STAD), testicular germ cell tumor (TGCT), thyroid carcinoma (THCA), thymoma (THYM), and uterine corpus endometrial carcinoma (UCEC) (Table 1 and Table S1). Eight datasets from six cancer types including MESO, PCPG, TGCT, THCA, THYM, and UCEC were excluded due to fewer than three survival datasets. Hence, our SurvivalML server included a total of 37,325 samples with both transcriptome data and survival information from 21 cancer types (Table 1 and Table S1).

**Figure 1.**
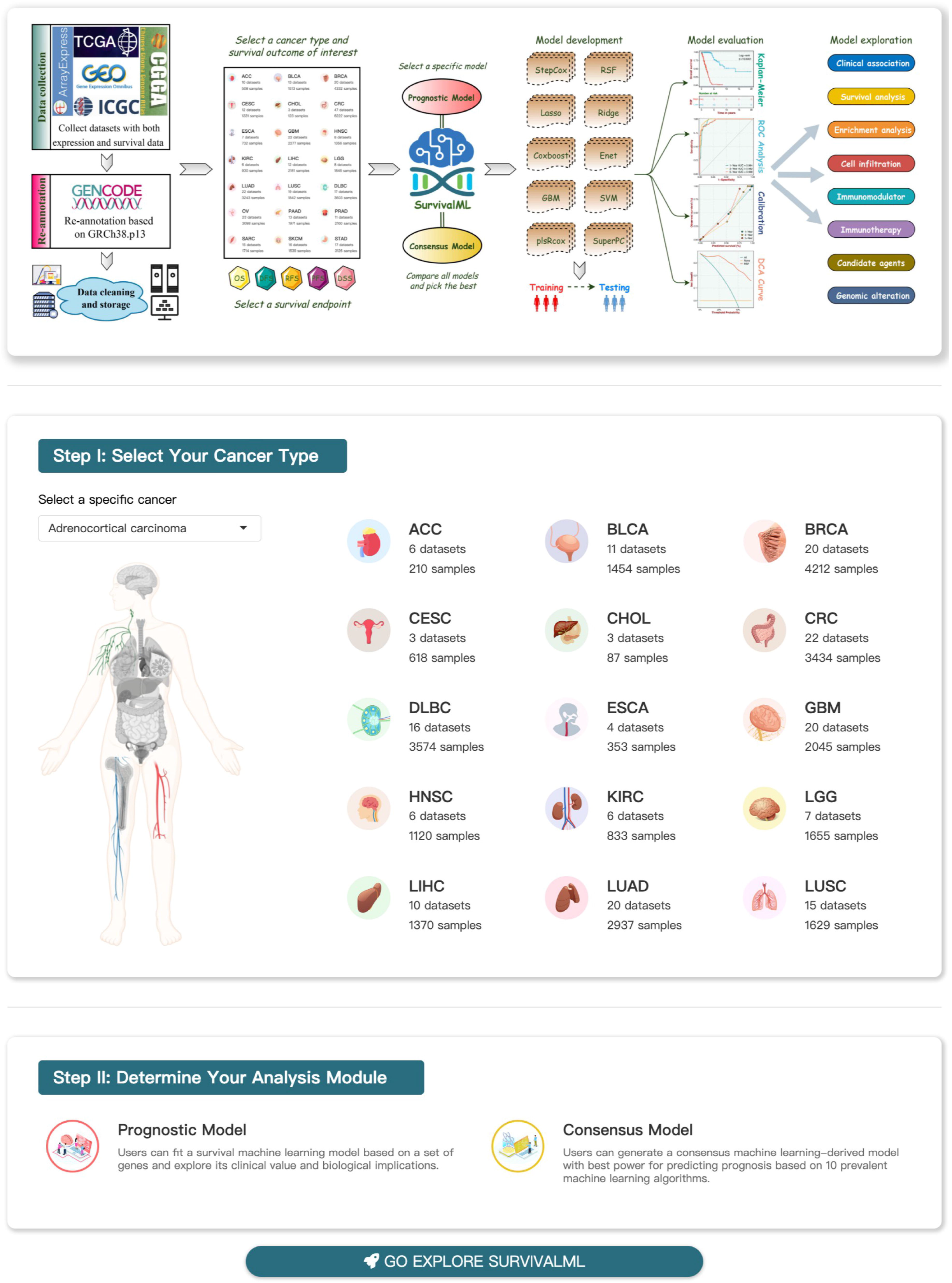
The home page of SurvivalML. The top panel indicates the workflow and main functions of SurvivalML. In Step I (the middle panel), users need to select an interesting cancer type from 21 solid tumors. In Step II (the bottom panel), they can determine an analysis module from two tabs, namely the prognostic model and the consensus model. The former module allows users to fit a survival model based on a specific set of genes. The latter module enables users to generate a consensus model with the best power for predicting prognosis according to 10 prevalent machine learning algorithms.

**Table 1.**
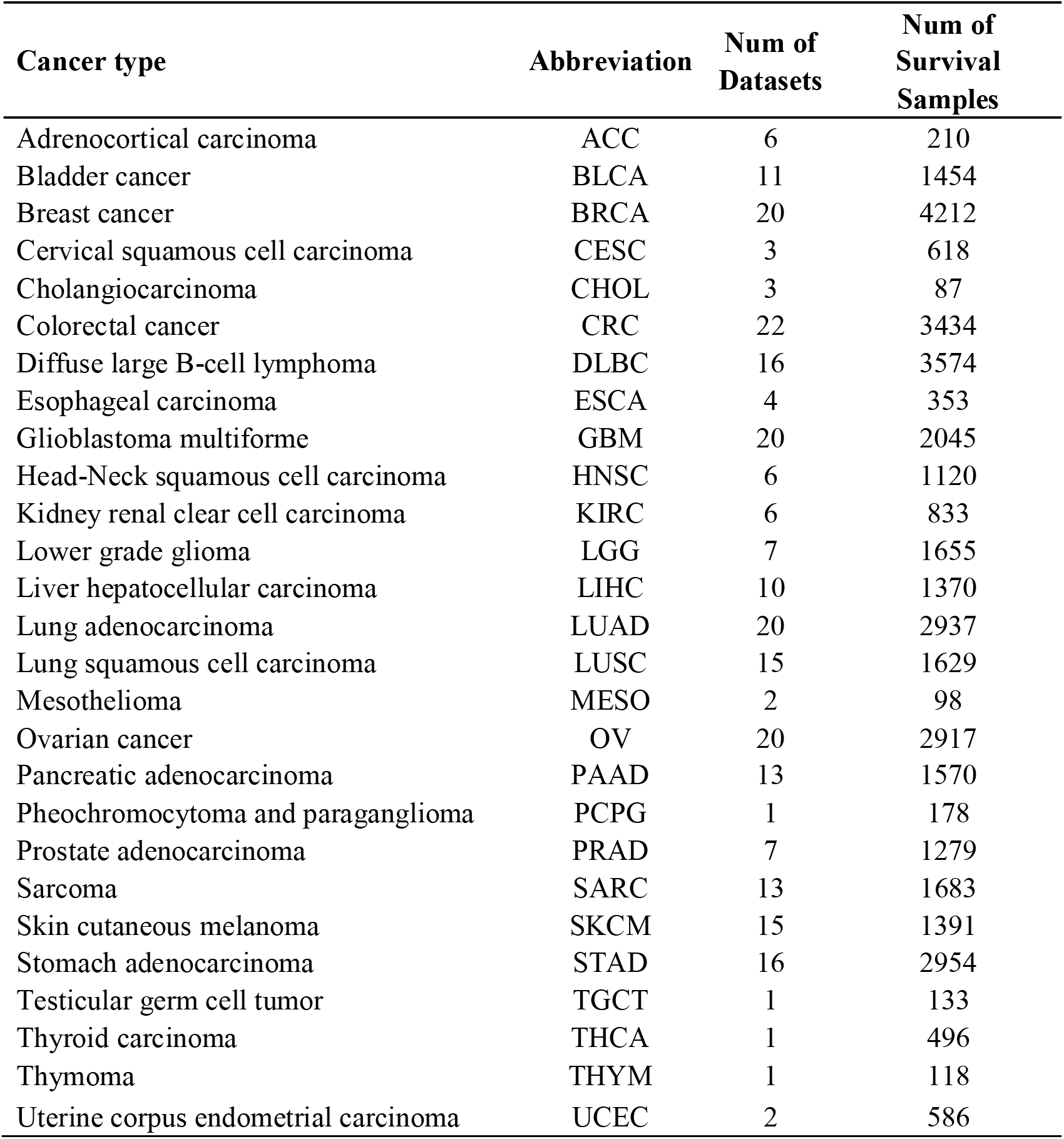
Dataset summary of 27 cancer types.

### Probe re-annotation

According to the GRCh38 patch 13 sequences reference from the GENCODE^17^ database (https://www.gencodegenes.org/), we re-annotated probesets using SeqMap^18^ if the original probe sequences FASTA files were available.

### Data processing

The workflow of data processing was as follows:

1. RNA-seq data (raw count or Fragments Per Kilobase Million) were uniformly converted to Transcripts Per Million values and further log transformed, which was more compatible with the distribution of microarray data^19^.
2. The chip raw data (e.g., CEL files) from Affymetrix^®^, Illumina^®^, and Agilent^®^ were processed using the *affy*^20^, *lumi*^21^, and *limma*^22^ packages, respectively. These packages encapsulated the functions of quality control and data normalization. Series normalization files were downloaded directly if microarray data were from other platforms, or the authors did not provide supplementary raw data.
3. The expression of each gene was transformed into a z-score in each dataset, which removed the effects of expression scale and dimension for model discovery^15^.

### Survival outcomes

SurvivalML enables users to develop and validate prognostic models for predicting five types of survival outcomes: overall survival (OS), disease-free survival (DFS), relapse-free survival (RFS), progression-free survival (PFS), and disease-specific survival (DSS) (Figure 1).

### Machine-learning algorithms

As previously reported^3^, this study retrieved 10 prevalent survival machine-learning algorithms, including stepwise Cox, random survival forest (RSF), least absolute shrinkage and selection operator (Lasso), Ridge, CoxBoost, elastic network (Enet), generalized boosted regression modeling (GBM), survival support vector machine (survival-SVM), partial least squares regression for Cox (plsRcox), and supervised principal components (SuperPC). Users can optimize the performance of survival models via flexibly tuning essential parameters and hyperparameters, which can be implemented interactively and conveniently on the SurvivalML website.

### Implementations

SurvivalML is a free online web application for helping the development and validation of survival models via various machine-learning algorithms, which was constructed by the Shiny app, HTML5, CSS, and JavaScript libraries. Processed data were stored on the server using R.data, allowing users to quickly recall data in the R environment. Users can interact with SurvivalML to further personalize their analytical needs. Analysis results are generally output in the form of images (PDF and PNG) or tables (CSV), which can be downloaded freely.

## Results

### Quick start

SurvivalML provides a simple interactive interface (Figure 1). Initially, users need to select an interesting cancer type from 21 solid tumors (Table 1). Subsequently, they can determine an analysis module from two tabs, namely the prognostic model and the consensus model (Figure 1). The former module allows users to fit a survival model based on a specific set of genes. The latter module enables users to generate a consensus model with the best power for predicting prognosis according to 10 prevalent machine learning algorithms. This study will delineate the operation processes and analysis results of these two modules.

### Prognostic model

#### Preparation

Taking lung adenocarcinoma (LUAD) as the cancer of interest, the preparation work of this module included six steps (Figure 2A):

1. Select an event outcome: Users needed to determine an event outcome of interest from five survival variables, including OS, RFS, DFS, PFS, and DSS. Here, OS was taken as our clinical endpoint.
2. Determine datasets with the selected outcome: Once the prognostic outcome was determined, this tab would display the available datasets with the selected outcome. Users could select several datasets for training and validation. Our database stored 21 datasets with OS information for LUAD. Here, we selected all datasets (Table S2).
3. Input your interest gene list: This tab required users to enter no less than two genes. Our server would detect whether the input genes were valid, in case the wrong gene ids were entered. Subsequently, users should define a name for their gene set, which tended to serve as the name of the prognostic score. Here, we entered 25 valid genes (Table S2) as well as customized them as “Pindex”. When everything was ready, clicked the submit button.
4. Gene summary: This tab summarized the distribution of the entered gene list across all selected datasets in the form of a bar graph and a text box. The bar graph illustrated the number of input genes presented in each dataset (Figure S1A) and the text box detailed which specific genes were missing from each dataset. This setting enabled users to flexibly increase or decrease specific gene features to control the number of included datasets for model training. Here, only 13 datasets had no absent genes (Table S2).
5. Split into training or validation datasets: Datasets with complete input genes were next assigned to training or validation datasets. Users could select a specific dataset as the training dataset, and the remaining datasets were utilized for validation. Here, we set TCGA-LUAD as the training dataset and the other 12 datasets were automatically grouped as the validation datasets (Table S2).
6. Choose a survival machine-learning algorithm: As mentioned above, we offered 10 algorithms for users to choose from. Users could determine a specific algorithm for model development in the training dataset. Here, random survival forest (RSF) was employed to train the survival model in TCGA-LUAD. After tuning parameters, users could click the red button of “Everything is ready! Start Modeling”. Notably, detailed explanations of all available parameters were illustrated on the Help page.

**Figure 2.**
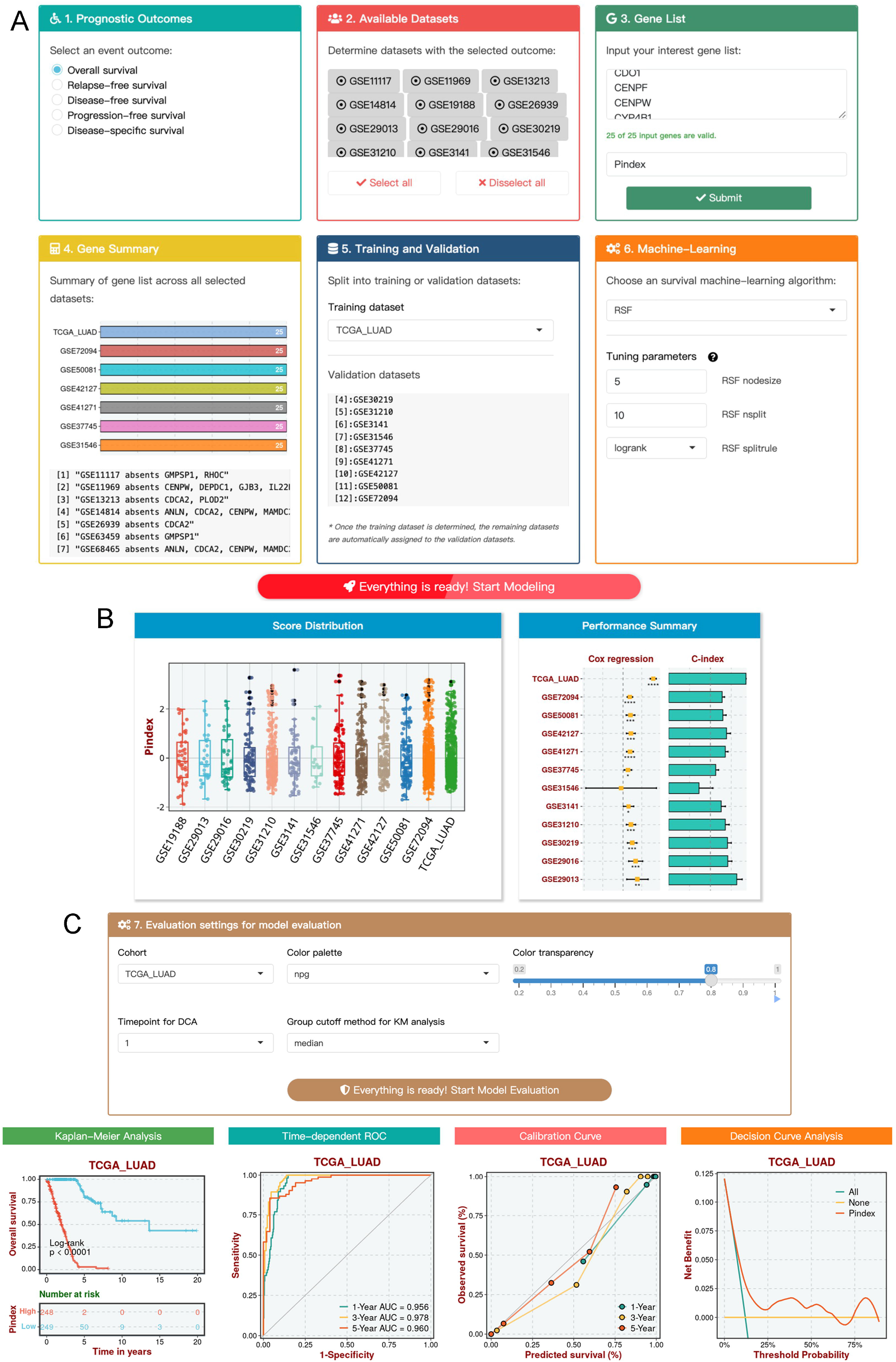
The operation processes and analysis results of the prognostic model. **A.** Taking lung adenocarcinoma as the cancer of interest, the preparation work of this module includes six steps: select an event outcome, determine datasets with the selected outcome, input your interest gene list, gene summary, split into training or validation datasets, and choose a survival machine-learning algorithm. **B.** The score distribution and performance summary of a model in all selected datasets. The box graph exhibits the score distribution of each dataset. Each dot represents a sample. The right panel presents Cox regression and C-index analysis results for the model across all datasets.

#### Model construction and evaluation

1. Score distribution and performance summary: Once the model construction was completed, a box graph would exhibit the score distribution of each dataset (Figure 2B). Each dot represented a sample. Additionally, Cox regression and C-index analysis results for the model across all datasets were generated (Figure 2B), thus enabling users to initially evaluate its performance. Detailed results of this example were illustrated in Figure S1B.
2. Model evaluation: The performance of a survival model was evaluated by four aspects: Kaplan-Meier survival analysis, time-dependent ROC, calibration curve, and decision curve analysis. Prior to model evaluation, users needed to select a dataset for evaluation, choose a color palette and transparency for Kaplan-Meier curve, and determine a timepoint for decision curve analysis (Figure 2C). Furthermore, users also should select a cutoff value for dichotomizing the prognostic score. This tab provided five cutoff methods, including median, mean, quantile, optimal, and custom. Here, we chose TCGA-LUAD to evaluate the performance of our Pindex signature. As shown in Figure 2C, Kaplan-Meier analysis revealed that patients with high Pindex possessed a significantly unfavorable OS relative to patients with low Pindex. Time-dependent ROC analysis assessed the discrimination of Pindex, with 1-, 3-, and 5-year AUCs of 0.956, 0.978, and 0.960 in TCGA-LUAD. Calibration curve was applied to compare predicted and observed survival probability, which demonstrated that Pindex had excellent calibration in predicting 1-, 3-, and 5-year OS. Decision curve analysis can be utilized to investigate whether our model outperformed “all” and “none” models. Here, Pindex exhibited better capacity in TCGA-LUAD. Of note, if users wanted to evaluate the performance of another dataset of the model, simply replaced the current dataset.

### Consensus model

#### Preparation

In this module, users did not need to provide a specific gene list, and gene features required for model training were jointly determined by all selected datasets. Here, we still chose LUAD as the cancer of interest. The preparation work of the consensus model also included six steps (Figure 3A):

1. Select an event outcome: Users could determine an event outcome of interest. In the example of this module, OS was also selected as our clinical endpoint.
2. Determine datasets with the selected outcome: As mentioned above, SurvivalML included 21 datasets with OS information for LUAD. Here, we selected 15 datasets for subsequent analysis (Table S3).
3. Dataset summary: After datasets were determined, SurvivalML would automatically keep only the intersection genes of all selected datasets. Here, 10,522 identical genes were expressed in 2,185 patients from 15 independent datasets.
4. Split into training or validation datasets: Users could determine a specific dataset as the training dataset, and the remaining datasets were utilized for validation. Here, we still assigned TCGA-LUAD as the training dataset and the other 14 datasets were automatically set as the validation datasets (Table S3).
5. Determine candidate genes: Our previous study systematically identified stably recurrence-related lncRNAs from multi-center cohorts and then developed a consensus machine-learning derived lncRNA signature that presented an excellent performance for predicting the recurrence risk of patients with colorectal cancer^23^. Hence, this tab enabled users to identify candidate genes that were stably linked to the survival outcome of interest via tuning the following three parameters: *P*-value for prognostic significance from Cox regression analysis, whether to screen out genes that were inconsistent in predicting prognosis (determined by hazard ratio) across statistically significant datasets, and the minimum number of statistically significant datasets. As shown in Figure 3A, we set Cox */*’-value cutoff to 0.1, genes with inconsistent prognosis to be filtered, and minimum number of significant cohorts to 9. After tuning parameters, the right panel would display a heatmap that revealed the associations between genes and prognosis across all datasets. White, red, and blue cells represented no significant, risky, and protective, respectively. In total, we generated 51 genes that were stably associated with OS in 15 LUAD datasets (Figure S2A).
6. Tune parameters for 10 algorithms: SurvivalML provided essential parameters and hyperparameters for 10 algorithms. Users could assess the detailed description of each available parameter on the Help page. Here, we utilized the default parameters, which were actually suitable for most cases. The purpose of developing 10 survival models was to identify the optimal one. In survival models, C-index is the most commonly used metric as it elegantly accounts for the censored data, and it is also the generalization of the ROC curve in survival data^24^. Compared with time-dependent ROC, C-index is a time-independent metric^25^. Thus, C-index can evaluate the overall discrimination of a survival model. In this module, the optimal model was determined via the mean C-index of all cohorts. Notably, a superior C-index of the training dataset due to model overfitting might heavily bias the mean C-index^3^, and we thereby provided an additional interactive setting: whether to include the performance of the training dataset to evaluate models. After accomplishing these settings, users could click the orange button of “Everything is ready! Start Modeling”.

**Figure 3.**
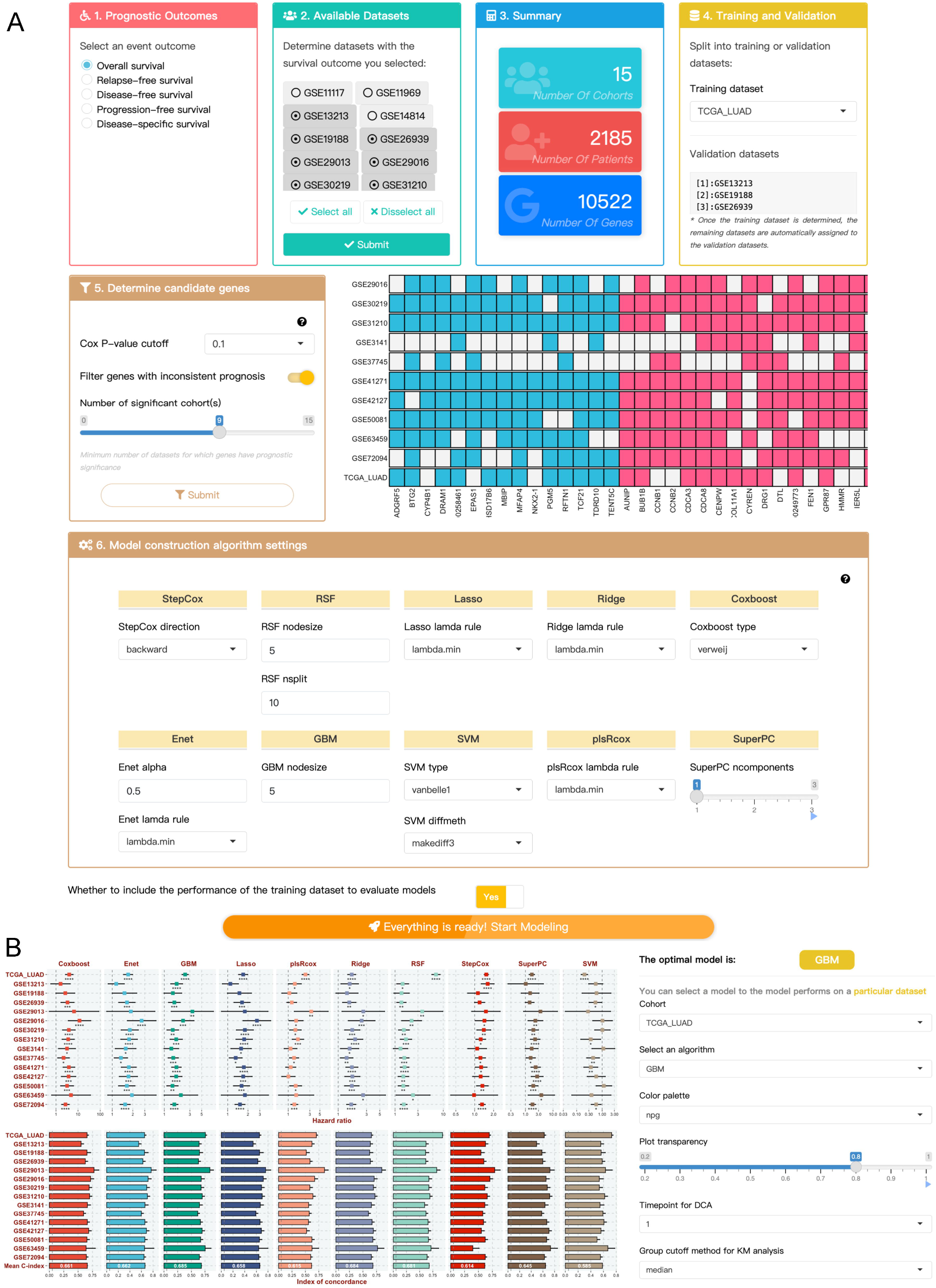
The operation processes and analysis results of the consensus model. **A.** The preparation work of the consensus model also includes six steps: select an event outcome, determine datasets with the selected outcome, dataset summary, split into training or validation datasets, determine candidate genes, and determine candidate genes. **B.** After model calculation, Cox regression and C-index analysis results for 10 models across all datasets are displayed. The optimal model with the highest mean C-index across all datasets is considered optimal. Here, the optimal one is generalized boosted regression modeling (GBM), which achieves a mean C-index of 0.685 across 15 independent datasets. Subsequently, users can select a specific cohort and tune personalized parameters to evaluate model performance.

#### Model construction and evaluation

1. Determine the optimal model: Model calculation of this module would take a relatively long time. Subsequently, Cox regression and C-index analysis results for 10 models across all datasets were displayed (Figure 3B). As mentioned above, the optimal model with the highest mean C-index across all datasets was considered the optimal one. Indeed, the model developed by this module possessed stronger generalization capabilities than most survival signatures, as previously reported^3,23,26–28^. Here, the optimal model was generalized boosted regression modeling (GBM), which achieved a mean C-index of 0.685 across 15 independent datasets (Figure 3B).
2. Evaluate model performance in each dataset: Users could select a specific cohort and tune personalized parameters to evaluate model performance (Figure 3B). In line with the prognostic model, this module also assessed the performance of the survival model via four aspects: Kaplan-Meier survival analysis, time-dependent ROC, calibration curve, and decision curve analysis (Figure S2B).

### Interface to the BEST web server

Our team developed a web server (BEST, https://rookieutopia.com/app_direct/BEST/) for comprehensive biomarker exploration on multi-center transcriptome data in solid tumors^29^. BEST enabled users to explore a single gene or gene list in according to the following eight aspects: clinical association, survival analysis, enrichment analysis, cell infiltration, immunomodulator, immunotherapy, candidate agents, and genomic alteration (Figure 1). SurvivalML focused on model development and validation, and each sample would generate a prognostic score based on model fitting, which was actually a continuous variable and could be analogous to gene expression. Thus, we retained the functions of the BEST website in the interface of SurvivalML, allowing users to use them directly for further exploration.

## Discussion

Cancer was endowed with aggressive biological characteristics and heterogeneous clinical outcomes. Clinicians routinely manage cancer patients with a focus on survival endpoints, generally including OS, DFS, RFS, PFS, and DSS. Thus, the ability to accurately predict prognosis and even identify high-risk patients that cannot be detected by conventional clinicopathological staging systems is the fundamental mission of prognostic models derived from gene expression profiles. However, the lack of rigorous validation in homogeneous multi-center datasets is a significant obstacle for current predictive models. A recent review demonstrated that although substantial studies established an association between gene signature and cancer prognosis, very few have been introduced into clinical settings due to a lack of reproducibility^8^. Hence, a platform dedicated to resolving the development and validation of survival models across multiple independent datasets is urgently imperative.

To the best of our knowledge, SurvivalML is the first and most comprehensive web server to date enabling users to flexibly develop and evaluate a survival model in multi-center datasets. SurvivalML provides large-scale data on 21 cancer types for model discovery. Currently, SurvivalML is also the largest and most comprehensive repository of RNA-seq/microarray cancer data with expression profiles and survival information. More importantly, these data were renewedly and uniformly re-annotated, normalized, and cleaned, which was more suitable for model validation among cross-platform cohorts. Following inclusion and exclusion criteria, SurvivalML included a total of 37,325 samples (253 datasets) with transcriptome data and survival information from 21 cancer types. All retained cancers harbored more than three independent datasets, ensuring multi-center validation. Hypothetically, if a gene signature maintains stable prognostic value across 20 BRCA datasets (n =4212) or 16 DLBC datasets (n =3574), then we have reason to believe that this signature can serve as a reliable and robust prognostic biomarker worthy of translation into clinical routine.

As an easy-to-use interactive web tool, SurvivalML allows users to conveniently develop and validate a survival model after determining a cancer type of interest. SurvivalML mainly contains two analysis modules, namely the prognostic model and the consensus model. The former module focuses on the prognostic significance of a specific (or customized) gene list in multiple datasets. Whereas the latter module is mainly applied to identify a stable homogeneous set of genes within selected multi-center datasets for modeling, and the optimal model is the one that harbors the best generalization ability (measured by C-index) among the 10 survival algorithms we provide. This is actually a response to questions raised by our previous research: why a specific algorithm should be employed, and which solution is the optimal one^23^. Numerous studies determined algorithms might lie primarily on the preference or bias of researchers. Therefore, it is crucial to provide multiple survival algorithms and matching parameter adjustments for those researchers without computational programming skills.

The performance evaluation of a model should be comprehensive, which can improve the tolerable error rate of the model in clinical feasibility. SurvivalML offers four aspects for the performance evaluation of survival models in each dataset, including Kaplan-Meier survival analysis, time-dependent ROC, calibration curve, and decision curve analysis, which are applied to assess their prognostic significance, discrimination, calibration, and latent clinical benefit, respectively. Users can assess these functions with great convenience as well as customize colors and approaches to generate personalized analysis results.

In addition, we provided an extra interface to the BEST web server, which was also developed by our team^29^. BEST enables users to explore the clinical significance and biological peculiarities of a single gene or gene list. Indeed, both gene expression and gene set scores were continuous variables, which are the same as the prognostic score generated from SurvivalML. Each patient corresponds to a specific value under all three patterns. Hence, users can further investigate other latent implications of the model, such as associations with clinicopathological traits, biological pathways, tumor microenvironment, immunotherapeutic efficacy, candidate agents, and genomic alteration. These functions derived from the BEST application can conduct a multi-dimensional exploration of a survival model, systematically exploiting the advantages of modern bioinformatics.

Although our SurvivalML web server is attractive, several limitations should also be acknowledged. First, the number of intersection genes in all datasets was relatively small. For example, only 11 genes overlapped in all bladder cancer datasets. As an alternative solution, users can remove datasets with fewer genes to generate more selection for modeling. Details of all datasets can be searched on the dataset page. Second, despite SurvivalML storing large-scale survival datasets, they were all retrospective and more common in sources of bias and confounding. Thus, survival models developed by SurvivalML need further validation in the prospective cohort.

To conclude, SurvivalML comprehensively retrieved and uniformly cleaned transcriptome data and survival information derived from 37,325 samples (253 datasets) of 21 cancer types, as well as provided an integrative framework for the development and validation of survival models across multiple independent datasets. Ten survival machine-learning algorithms and rounded approaches for evaluating model performance were offered to establish a survival model flexibly and systematically. An extra interface to the BEST web server allowed users to further explore the clinical significance and biological peculiarities of a model. With constant user feedback and further improvement, SurvivalML can serve as a promising platform to develop cancer survival models in multi-center datasets.

## Data availability

R studio is a language and environment for statistical computing, which is available at https://www.r-project.org/. Shiny is an open-source collaborative initiative, which is available in the GitHub repository (https://github.com/rstudio/shiny). SurvivalML (https://rookieutopia.com/app_direct/SurvivalML/) is free and open to all users, with no login requirements. Data were collected from The Cancer Genome Atlas Program (TCGA, https://portal.gdc.cancer.gov), Gene Expression Omnibus (GEO, https://www.ncbi.nlm.nih.gov/geo/), International Cancer Genome Consortium (ICGC, https://dcc.icgc.org), Chinese Glioma Genome Atlas (CGGA, http://www.cgga.org.cn/), and ArrayExpress (https://www.ebi.ac.uk/arrayexpress/). The processed data used in this study are available from the corresponding authors upon request.

## Acknowledgments

Not applicable.

## Author contributions

ZQL designed and developed this web server. XWH provided personal fund support and supervised this project. ZQL, LL, HX, SYW, ZX, XYG, LBW, and CGG collected and processed essential data. ZQL wrote and edited this manuscript. SC, QC, PL, and JZ revised this manuscript. All authors approved this manuscript.

## Ethics approval and consent to participate

Not applicable.

---

**Figure S1. Gene summary of each dataset and performance summary across all datasets. A.** The bar graph illustrated the number of input genes presented in each dataset. **B.** Cox regression and C-index analysis results for the model across all datasets.

**Figure S2. Generation of stable prognosis-related genes and model evaluation. A.** A total of 51 genes were stably associated with overall survival in 15 lung adenocarcinoma datasets. **B.** The performance of the survival model was evaluated via four aspects: Kaplan-Meier survival analysis, time-dependent ROC, calibration curve, and decision curve analysis.

