## Supplementary figures and images for "SurvivalML: an integrative platform for the discovery and exploration of prognostic models in multi-center cancer cohorts"

### Figure S1

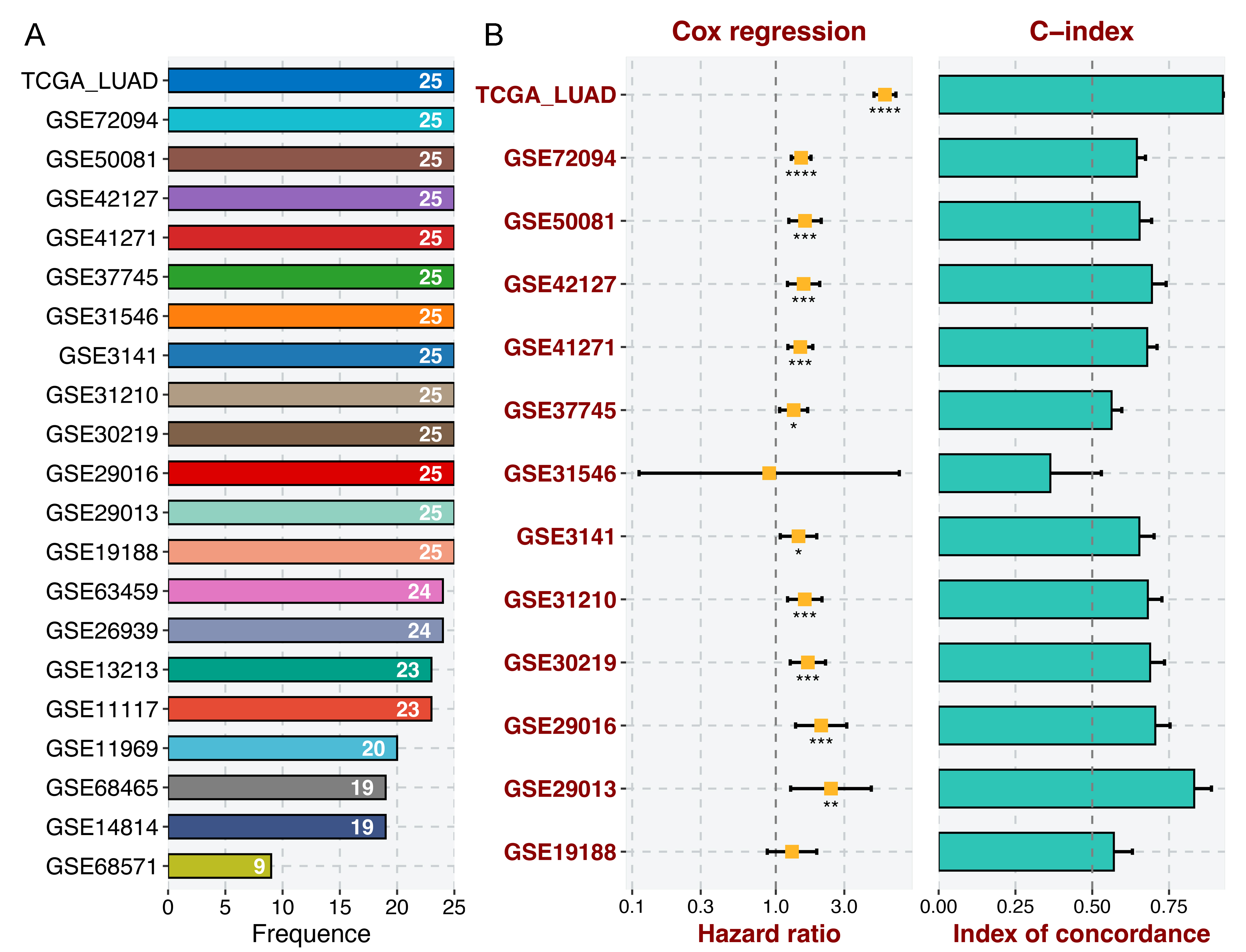

### Figure S2

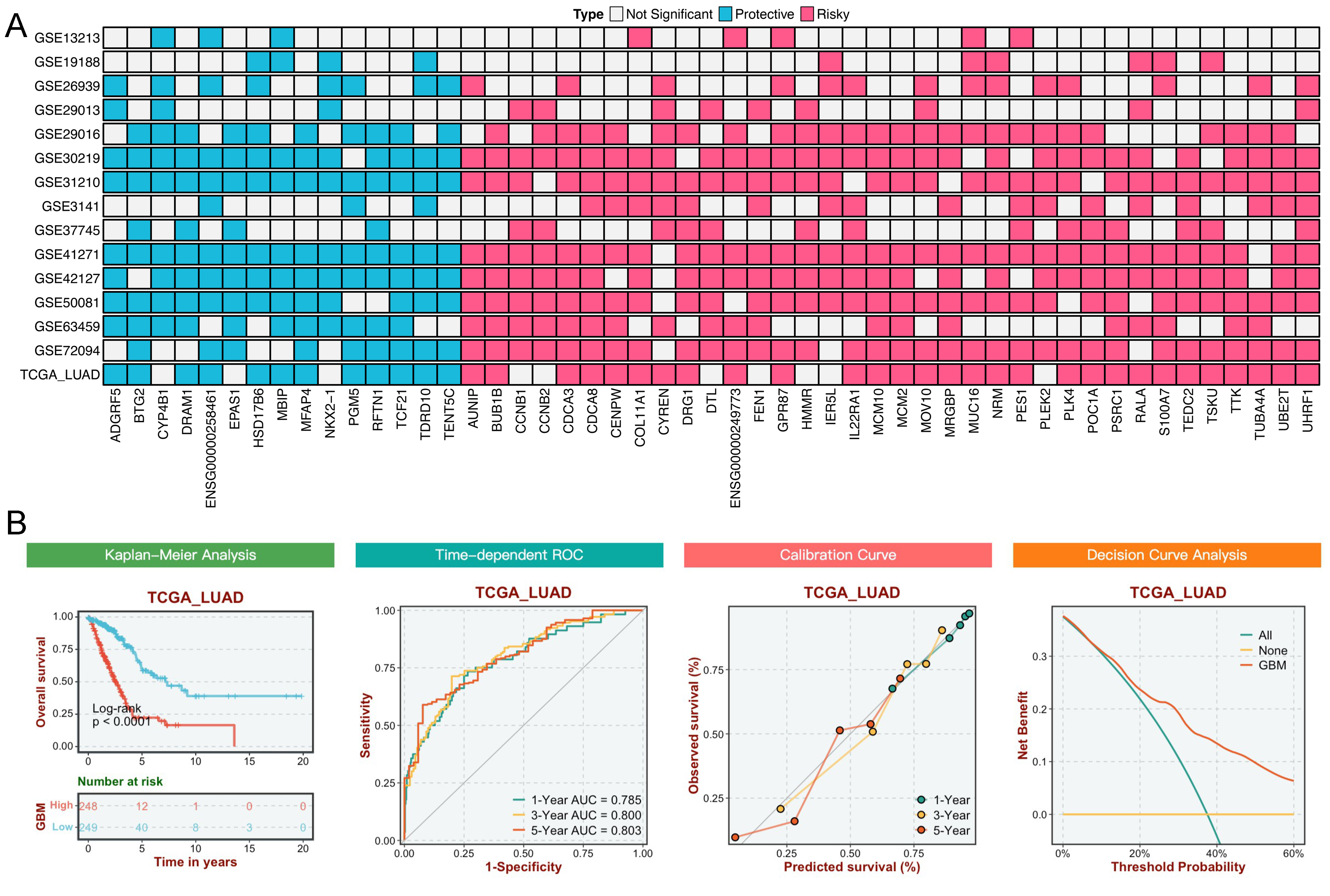
